# Cryo-EM structure of a conjugative T4SS identifies a molecular switch regulating pilus biogenesis

**DOI:** 10.1101/2024.02.08.579498

**Authors:** Kévin Macé, Gabriel Waksman

## Abstract

Conjugative Type IV Secretion Systems (T4SS) mediate bacterial conjugation, a process that enables the unidirectional exchange of genetic materials between a donor and a recipient bacterial cell. Bacterial conjugation is the primary means by which antibiotic resistance genes spread among bacterial populations (Barlow, 2009; Virolle *et al*, 2020). Conjugative T4SSs elaborate a long extracellular filament, termed “pilus” to connect with the recipient cell. Previously, we solved the cryo-EM structure of a conjugative T4SS. In this article, based on additional data, we present a more complete T4SS cryo-EM structure than that published earlier. Novel structural features include details of the mismatch symmetry within the OMCC, the presence of a 4^th^ VirB8 subunit in the asymmetric unit of both the Arches and the IMC, and a hydrophobic VirB5 tip in the distal end of the stalk. However, more significantly, we provide previously-undescribed structural insights into the protein VirB10 and identify a novel regulation mechanism of T4SS-mediated pilus biogenesis by this protein, that we believe is a key checkpoint for this process.

## INTRODUCTION

Conjugative Type IV Secretion Systems (T4SSs) minimally contain 12 proteins, named VirB1-VirB11 and VirD4 (Costa *et al*, 2020; Costa *et al*, 2023; Waksman, 2019). VirB1 is a lytic transglycosylase, facilitating T4SS assembly (Chandran Darbari & Waksman, 2015). VirB2 is the pilin subunit located into the inner membrane (IM), which is subsequently extracted from the IM and assembled to form a helical extracellular appendage termed “the conjugative pilus” (Costa *et al*, 2016). The recent cryo-Electron Microscopy (cryo-EM) structure of the conjugative Type 4 Secretion System (T4SS) encoded by the plasmid R388 revealed that the remaining VirB proteins form a large assembly organised into four subcomplexes (Figure 1): the Outer Membrane Core Complex (OMCC), the Stalk, the Arches and the Inner Membrane Complex (IMC) (Mace *et al*, 2022). The OMCC is made of TrwH/VirB7, TrwF/VirB9, and TrwE/VirB10 (in this study, we use both the R388 Trw and *Agrobacterium* VirB naming nomenclatures). It is itself composed of two parts: the 14-fold symmetrical O-layer embedded within the outer membrane (OM) and the 16-fold symmetrical I-layer beneath it. Previous structural studies of the OMCC have revealed that the O-layer and I-layer often present a mismatch symmetry (Amin *et al*, 2021; Durie *et al*, 2020; Sheedlo *et al*, 2020), and yet they are both composed of the same proteins (for example, TrwF/VirB9 and TrwE/VirB10). The Stalk is 5-fold symmetrical and is made of TrwJ/VirB5 and TrwI/VirB6, with TrwI/VirB6 embedded into the IM and hypothesized to serve as a platform for pilus subunit recruitment and assembly (Mace *et al*, 2022). Indeed, TrwI/VirB6 features two key sites: a VirB2 binding site at its base in the IM, and a pilus-assembly site at its top. TrwJ/VirB5 is located at the tip of the Stalk, and is expected to be recruited at the pilus tip during the early step of pilus biogenesis. The 6-fold symmetrical Arches are observed in the previously published structure to be made of 6 trimers of the periplasmic domain of TrwG/VirB8 (TrwG/VirB8_peri_) forming a large ring around the Stalk. The 6-fold symmetrical IMC is made of 6 copies of a protomer comprising 1 copy of TrwM/VirB3, 2 copies of the TrwK/VirB4 ATPase subunits termed TrwK/VirB4_central_ and TrwK/VirB4_outside_, and in the structure published previously three N-terminal tails of TrwG/VirB8 (TrwG/VirB8_tail_). Six TrwK/VirB4_central_ form a central hexamer linked to the IM by VirB3. TrwK/VirB4_outside_ forms a robust dimer with VirB4_central_ and interact with the TrwG/VirB8_tail_. TrwD/VirB11 was not part of the complex because detergent solubilisation of the complex results in the dissociation of this ATPase. However, its location under the TrwK/VirB4_central_ hexamer was ascertained by co-evolution and site-directed mutagenesis (Mace *et al*., 2022). Although a third ATPase, TrwC/VirD4, is known to be part of the IMC at some point during conjugation, no density could be found for it and therefore its location and interactions with other T4SS components remain elusive.

**Figure 1.**
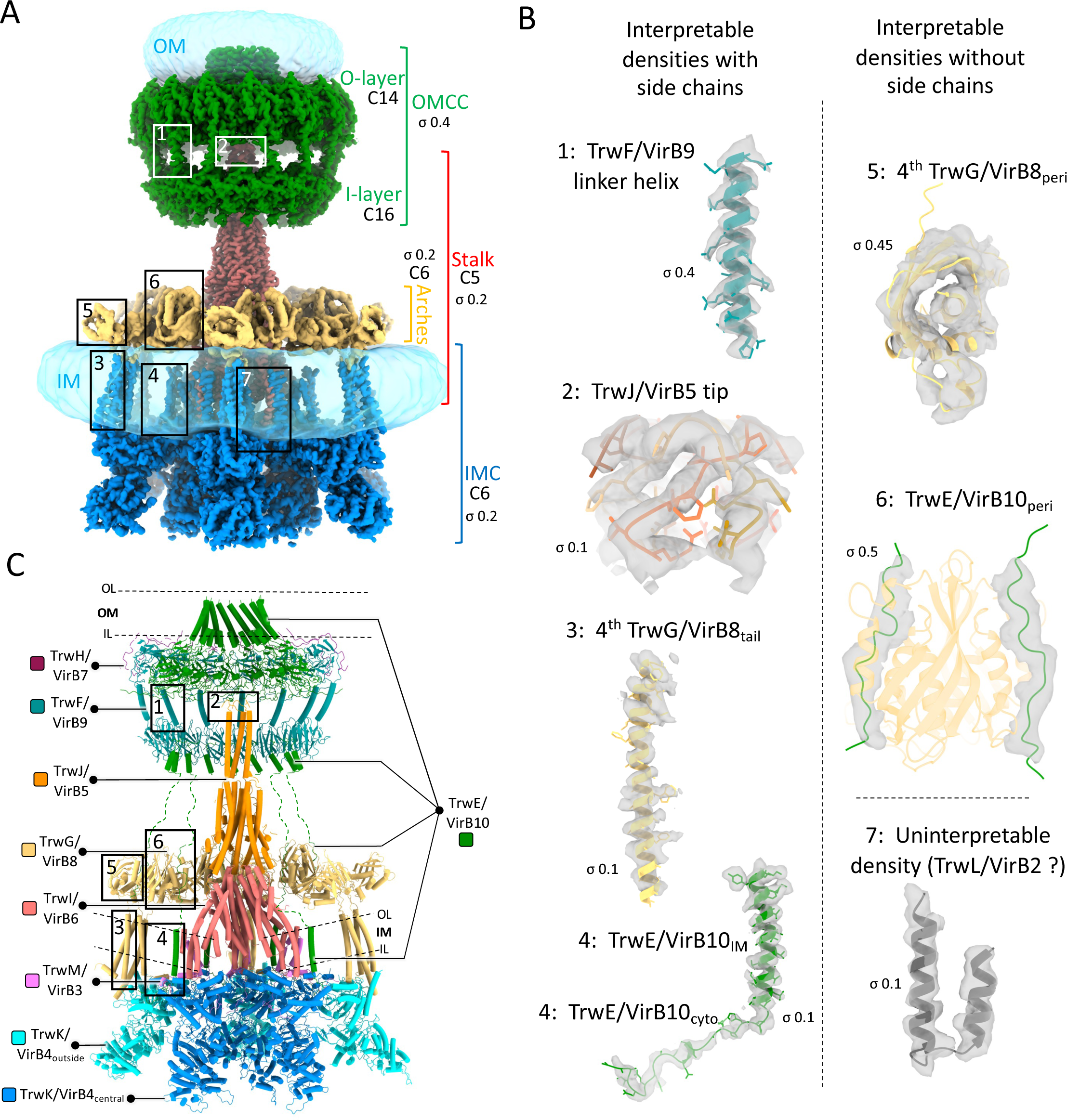
Improved structure of the R388 conjugative T4SS. **A.** Composite electron density map. A composite electron density map of the R388 T4SS is presented, created by assembling maps detailed in Table EV1A. In this map, sub-complexes of the T4SS, including the Outer Membrane Core Complex (OMCC; map referred to as “OMCC Conformation A at 3.2 Å”), the Stalk (map referred to as “Stalk C5 at 3.0 Å”), the Arches (map referred to as “Arches at 6.2 Å”), and the Inner Membrane Complex (IMC, map referred to as “Extended IMC protomer at 3.8 Å”), are color-coded in green, red, yellow, and blue, respectively. Sigma levels for each map is reported. Symmetry within the sub-complexes is indicated. The detergent and/or lipid densities at the membrane and outer membranes are depicted as semi-transparent light blue density. Additionally, newly identified densities from this study compared to the one published previously are highlighting in rectangular boxes and detailed in panel B. **B.** Newly identified densities. Three categories of newly identified densities are shown as grey, semi-transparent surface contour: interpretable with side chains (regions 1 to 4), interpretable with only secondary structures represented (no side chains; regions 5 and 6), and uninterpretable (region 7). Structures shown in these densities are in cartoon representation, with side chains reported only for regions 1 to 4. The structures are color-coded according to proteins as in panel C. Sigma levels are reported. For region 7, the density was tentatively ascribed to TrwL/VirB2 but in the absence of corroborating evidence, the density is classed as uninterpretable at this moment in time. **C.** Composite model of the T4SS. A composite model of the R388 T4SS is presented in cartoon representation colour-coded per proteins as shown in margins. Positions of IM and OM based on cryo-EM densities of detergents and lipids are shown by dashed lines. Regions highlighted in B are shown in correspondingly-numbered boxes.

In the cryo-EM map published previously, certain regions of the complex still presented challenges due to insufficient resolution, preventing a detailed interpretation of some part of the electron density map. In response to these limitations, we present here an enhanced T4SS structure that builds upon our previous work (Table EV1, Figures EV1 and EV2). This improved structure was achieved by collecting additional cryo-EM data as well as dedicated image processing on regions of interest. Specifically, we generated a total of 9 cryo-EM maps, with a range of resolutions of 2.5 Å to 6.2 Å, to gain additional structural insights. These include the elucidation of several conformations of the OMCC, the structure and interactions of the TrwE/VirB10 N-terminus comprising periplasmic, inner membrane, and cytoplasmic regions, the tip region of TrwJ/VirB5, a fourth TrwG/VirB8 subunit in the asymmetric unit of the Arches and IMC, and the connector domain of the Arches formed by the collective contribution of all four TrwG/VirB8 subunits. All structures are validated by co-evolution and AlphaFold modelling. Altogether, our results greatly enhance our understanding of T4SS molecular mechanisms, notably its regulation of pilus biogenesis by TrwE/VirB10.

## RESULTS AND DISCUSSION

In the higher resolution cryo-EM maps presented here, additional densities were observed, that fall into 3 categories: i-densities with clear secondary structural features and side chains, where accurate models could be built *de novo* (regions 1 to 4 in Figure 1), ii-densities with clear secondary structural elements but no side chains, where the main chain could be docked accurately, but no side chains could be built (regions 5 and 6 in Figure 1) and finally iii-insufficiently resolved densities which could not be interpreted (region 7 in Figure 1). The latter showed 2 tubes of densities in the IM that could possibly correspond to TrwL/VirB2 known to contain 2 TMs, but this interpretation could not be validated and therefore will not be discussed further here.

### The outer membrane core complex (OMCC)

All resolved OMCC structures consistently exhibit a conserved global organisation composed of an O-layer and I-layer sub-complex, often displaying a mismatched symmetry between them (Amin *et al*., 2021; Durie *et al*., 2020; Sheedlo *et al*., 2020). For instance, the OMCC of R388 has a C14/C16 mismatch between the O- and I-layer, respectively (Mace *et al*., 2022) (Figure 2). Intriguingly, O- and I-layers are formed of the same proteins: to only mention R388, the O-layer is formed of the CTD of both TrwF/virB9 and TrwE/VirB10, while the I-layer is formed of the NTD of these same proteins. Unsymmetrised C1 maps for the entire OMCCs are obtainable, however, they are often of insufficient resolution to derive important structural features such as the TrwF/VirB9 linker sequence between the two layers or to derive detailed information on the symmetry mismatch. In this study, thanks to a larger data set, we were able to achieve high resolution without imposing any symmetry constraints (Figure EV1), enabling a more in-depth investigation of the OMCC.

**Figure 2.**
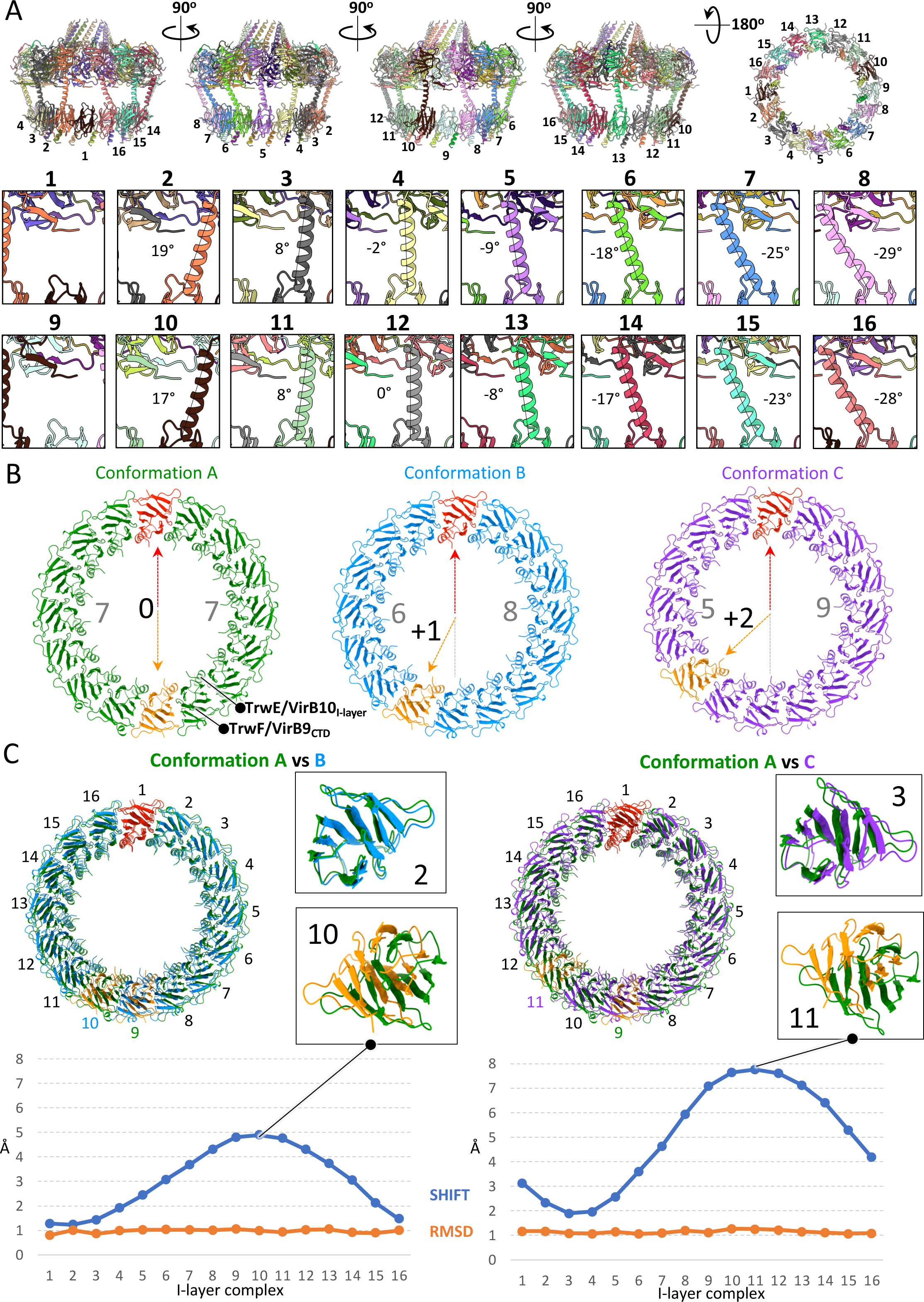
Analysis of OMCC mismatch symmetry. **A.** Structure analysis of the OMCC conformation A. In the top panel, five views of the OMCC structure obtained without symmetry applied (from the “Conformation A at 3.2 Å” map) are shown in cartoon representation and coloured by chains. Each I-layer complex is numbered. The bottom panel provides a close-up view of all TrwF/VirB9 linker helices connecting the O-layer to the I-layer, with the angles of the helices relative to the vertical axis indicated. **B.** Three distinct conformations of the OMCC. In this panel, three OMCC structures are presented, each exhibiting a distinct I-layer organisation. A top view of the three I-layer structures is shown in cartoon representation, with colours representing different conformations: green for the conformation A, blue for the conformation B, and purple for the conformation C. For each conformation, the two extra I-layer complexes are coloured in red and orange. The number of I-layer complexes between the two inserted extra I-layer complexes is indicated in grey. **C.** Superposition of OMCC structures. This panel shows pairwise superpositions of the OMCC structures presented in panel B, with alignment based on the O-layer. Left: superposition of Conformations A and B. Right: superposition of Conformations A and C. The impact of the insertion of two extra complexes in the I-layer is shown at right of each superposition in zoom-in panels illustrating the least and most affected complexes (2 and 10 for Conformation A versus B and 3 and 11 for Conformation A versus C). Also reported underneath is the shift (as reported using Chimera) separating the two superposed structures for each I-layer complex position. Despite these shifts, each I-layer complex structures remains identical, as evidenced by the RSMD values (orange line).

Initially, we solved the OMCC structure without imposing symmetry at 3.1 Å resolution (Figures 2A and EV1). In this map, the two additional I-layer binary complexes of TrwF/VirB9_NTD_ and TrwE/VirB10_NTD_ are inserted in diametrically opposite locations (Figure 2A). Clear density is seen for 14 linker sequences (residues 128 to 150) between the two domains of TrwF/VirB9, which we can now confirm is mostly α-helical. As anticipated, the biggest impact of domains insertion in the I-layer is observed in this linker helix. When the angle it makes relative to a vertical axis is measured, a pattern can be seen, with the largest angles observed on each side of the insertion (see complexes 2 and 10 or 8 and 16 in Figure 2A, lower panels). The angle tappers off the further the complex is from both insertions (see complexes 4 and 12 in the same Figure panels). Clearly, linker helix flexibility is crucial to accommodate complex insertion in the I-layer.

Further analysis employing 3D classification (Figure EV1) reveals a degree of heterogeneity in the positioning of the two extra TrwF/VirB9_NTD_-TrwE/VirB10_NTD_ sub-complexes within the I-layer (Figure 2B). We obtained three distinct OMCC structures, each characterised by a unique arrangement of the two extra sub-complexes in the I-layer. These three arrangements are: i) “conformation A” which is the one observed previously (see above) with the extra sub-complexes inserted diametrically opposite of each other; ii) “conformation B” where insertion of the second extra sub-complex in the I-layer is shifted by 1 compared to conformation A; in this conformation, there are 6 and 8 TrwF/VirB9_NTD_-TrwE/VirB10_NTD_ sub-complexes on each side of the extra sub-complexes; iii) “conformation C” where insertion of the second extra sub-complex is shifted by 2 compared to conformation A; in this conformation, 5 and 9 TrwF/VirB9_NTD_-TrwE/VirB10_NTD_ complexes are observed on each side of the extra sub-complexes. Aligning the structures of these three I-layer conformations using the O-layer as the superimposing entity underscores how the variability in the positions of the two extra I-layer complexes impacts the overall I-layer structure (Figure 2C). Although the structure of each complexes forming the I-layer do not change significantly (RMSD of 1.06 Å), a shift between corresponding complexes in the superposition of conformations A and B or A and C is observed. In both, this shift changes in magnitude, being maximal in the superposition involving the second extra sub-complex (that in orange in Figure 2C i.e. complex 10 in the A versus B superposition and complex 11 in the A versus C superposition) and, intriguingly, being minimal in the complexes opposite (complex 2 in the A versus B superposition and complex 3 in the A versus C superposition). Thus, insertion of extra complexes occurs at various positions in the I-layer, resulting in structural adjustments that propagate over the entire I-layer. The conservation of this mismatch symmetry across various T4SS types emphasizes its importance. This distinct pattern observed in the OMCC, characterized by a quasi-symmetrical mismatch, suggests the existence of a specific maturation process during OMCC assembly, although the exact mechanism remains enigmatic. The authors suggest a plausible role for this asymmetry: to expand the dimensions of the I-layer structure, thereby achieving the ideal diameter to accommodate the growing pilus at a later stage.

### The Stalk and Arches

In these regions of the T4SS structure, improved resolution resulted in better defined densities at the tip of TrwJ/VirB5. Also, a 4^th^ subunit of TrwG/VirB8 in the asymmetric unit of both the Arches (TrwG/VirB8_peri_) and the IMC (TrwG/VirB8_tail_) could be placed and built.

In the TrwJ/VirB5 N-terminal tip (Figure 3A), 11 residues could be added at the N-terminus (residues Gln23-Ala33). Gln23 is the very N-terminal residue in the protein after signal sequence cleavage. As shown in Figure 3A, middle panel, this newly built region is hydrophobic in nature. VirB5 homologues have been shown to locate at the tip of the pilus, an ideal location to interact with the recipient cell surface (Aly & Baron, 2007). Viewed from the top, the hydrophobic tip of TrwJ/VirB5 resembles a needle (Figure 3A, insets of each panel). It is not known whether VirB5 proteins can penetrate membranes. However, a body of structural work on this family of proteins ranging from their structural similarities with hemolysin E (Mace *et al*., 2022) and the hydrophobic nature of their N-terminal tip appear to point to a role in recipient membrane recognition and possibly puncturing.

**Figure 3.**
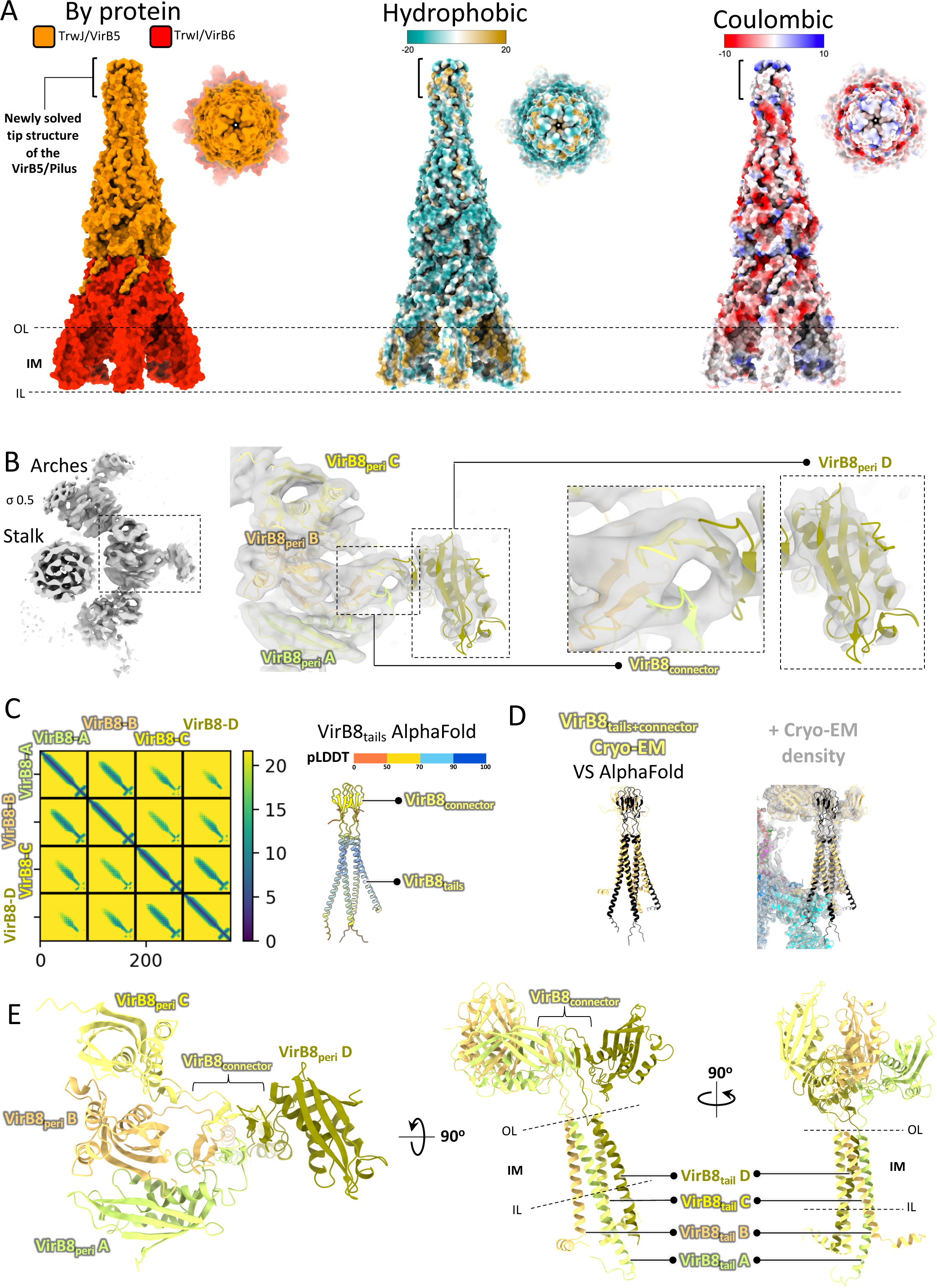
Structure of the Stalk and Arches subcomplex. **A.** The stalk in various representations. In this panel, the Stalk structure is presented in both side and top views, employing three different colour-coding schemes: by proteins (left), hydrophobicity (middle), and coulombic properties (right). The newly resolved tip of the TrwJ/VirB5 structures is indicated in brackets, and the IM is delineated by two dashed lines. **B.** Details of the cryo-EM map in the Arches region. Left: top view of the “Arches at 6.2 Å” map (Table EV1a) is displayed in grey colour and shows three asymmetric units, one of them (in dashed line box) is better defined. Sigma level is indicated. Middle panel: close-up of the asymmetric unit of the “Arches at 6.2 Å” map corresponding to the region shown in inset in left panel. The structure of four TrwG/VirB8_peri_ and four TrwG/VirB8_connector_ regions are shown in cartoon representation (rigid body fitting correlation coefficient of 0.88 and 0.75 for TrwG/VirB8_peri_ and TrwG/VirB8_connector_, respectively). Density is shown in semi-transparent grey. Two dashed rectangles highlight two newly solved portions of the structure compared to our earlier work: the TrwG/VirB8_connector_ and the 4^th^ TrwG/VirB8_peri_, which are further detailed in the zoomed-in views on the right. **C.** TrwG/VirB8 Arches structure validation using AlphaFold. Co-evolution plot (left) and AlphaFold structure colour-coded by model quality (pLDDT; right) for the four TrwG/VirB8 tail and connector domains. Coevolution residue pairs detected by AlphaFold are marked with green dots on the diagram at left. **D.** Right panels: superposition of the AlphaFold (in black ribbon) and cryo-EM (in yellow ribbon) models for the TrwG/VirB8 connector and tail domains (RMSD of 9.398 and 6.928 Å for TrwG/VirB8_connector_ and TrwG/VirB8_tails_, respectively) without (left) or with (right) the cryo-EM electron density map for this region contoured at same sigma level as in panel B. **E.** Structure of the asymmetric unit of the TrwG/VirB8 Arches structure. The four TrwG/VirB8 subunits are color-coded in four shades of yellow and in cartoon representation. TrwG/VirB8 subunits and domains are indicated, as well as the location of the IM represented by two dashed lines.

In our previous report (Mace *et al*., 2022), resolution for the Arches was low due to the considerable flexibility of the region. Thus, only three TrwG/VirB8 periplasmic domains (TrwG/VirB8_peri_) in the asymmetric unit (18 total) could be located unambiguously. However, density was observed on the side of the TrwG/VirB8_peri_ ring, density that could not be interpreted at the time. Additional data collection has however resulted here in improved density in this region, allowing to ascribe it unambiguously to a 4^th^ TrwG/VirB8_peri_ domain. As shown in Figure 3B, this density is shaped like a hook and protrudes outwards away from the Arches ring. In the hook, one molecule of TrwG/VirB8_peri_ could be fitted based on secondary structures (see Figure 1B, region number 5, and Figure 3B, right most panel). Resolution was however too low to build side chains in this region. Correspondingly, a 4^th^ TrwG/VirB8_tail_ is observed in the asymmetric unit of the IMC (Figures 3C and 3D). In this region of the IMC, side chains are clearly visible and a complete model of the 4^th^ TrwG/VirB8_tail_ was built. AlphaFold (Jumper *et al*, 2021) also predicts a four-helix bundle between the four TrwG/VirB8_tail_ and, reassuringly, this AlphaFold model superimpose well with the one built in the improved density (Figures 3C and 3D). Therefore, overall, there are 24 TrwG/VirB8 subunits in the T4SS.

The three TrwG/VirB8_peri_ domains in the asymmetric unit of the Arches described previously were related to each other in a way seen in two different earlier publications: MolA and MolB forms an interface similar to that observed in *Helicobacter pylori* CagV/VirB8_peri_ while the MolB and MolC interface is similar to that observed in *Brucella suis* VirB8_peri_ (Terradot *et al*, 2005; Wu *et al*, 2019). The newly observed MolD makes only very few contacts with this trimeric unit and is not related by symmetry with any of the other 3 subunits. However, it appears stabilised (and therefore visible in the density) mostly via interactions of its connector sequence (between TrwG/VirB8_peri_ and TrwG/VirB8_tail_, residues 63 to 94) with the corresponding connector sequences of the other 3 subunits (Figure 3, C-E). Each connector sequence forms a β-hairpin, the four of them assembling to form a 8-stranded β-sandwich (Figure 3, C-E).

Unexpectedly, two additional densities were observed running along the long axis of TrwG/VirB8_peri_ MolB and MolC (in green in Figure 4A). These densities showed no side chains and therefore could not be unambiguously assigned. Nevertheless, co-evolution analysis and AlphaFold modeling (Figure 4B) led us to hypothesize that these densities correspond to the sequence of TrwE/VirB10 between residues 83 and 101, a region we label TrwE/VirB10_Arches_. This region interacts with residues in the α-helical side of TrwG/VirB8_peri_ (α4-α6) (Figure 4C). Previous studies have described unidentified sequences of VirB10 to interact with VirB8_peri_ in the α4-α5 and the β1 regions (Sharifahmadian *et al*, 2017). In the structure of the fully-assembled T4SS presented here and earlier (Mace *et al*., 2022), the β1 region is implicated in intra- and inter-asymmetric unit interactions (Figure 4C). Therefore β1 is unavailable for binding to VirB10. However, as shown in Figure 4D in red, the α4-α5 region mapped previously (Sharifahmadian *et al*, 2017) to interact with VirB10 is exactly where we observed the TrwE/VirB10_Arches_ density. Therefore, it is indeed likely that the densities observed belong to TrwEVirB10 and the assignment based on AlphaFold and co-evolution is correct. No TrwE/VirB10_Arches_ sequences were found bound to TrwG/VirB8_peri_ MolA or MolD. Interestingly, although there are 16 copies of the TrwE/VirB10_Arches_ sequence available in the T4SS structure for binding to VirB8_peri_, only 12 would bind to the VirB8_peri_ Arches region (2 per asymmetric unit in a region that is 6-fold symmetrical).

**Figure 4.**
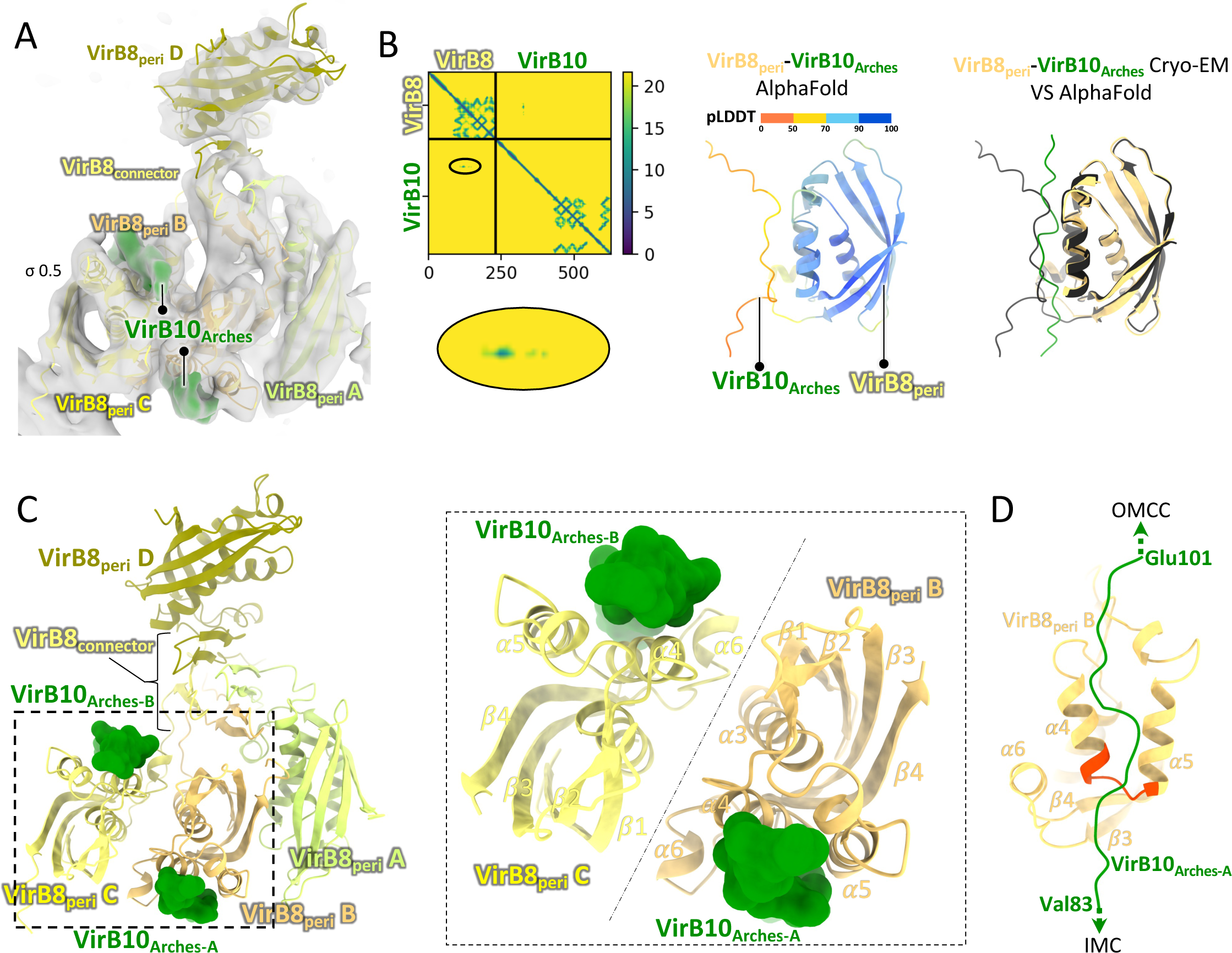
Structure of the Arches subcomplex and VirB8-VirB10 interaction. **A.** Top view of the “Arches at 6.2 Å” cryo-EM map showing the two densities corresponding to TrwE/VirB10_Arches_. The map is presented in semi-transparent grey (sigma level indicated). The four TrwG/VirB8 are fitted within this map as represented in Figure 3, panel E. Two densities corresponding to VirB10_Arches_ are coloured in green. **B.** Identification TrwE/VirB10_Arches_, the region of TrwE/VirB10 that interacts with TrwG/VirB8_peri_ using co-evolution and AlphaFold. Left: co-evolution analysis between VirB10 and VirB8, with co-evolving residue pairs shown as green dots surrounded by a solid-lined oval. A zoom-up of this region is shown underneath. Middle: AlphaFold-derived structural model of TrwG/VirB8_peri_-TrwE/VirB10_Arches_ (residues 88 to 96) shown in cartoon representation, colour-coded by model quality (pLDDT). Right: superposition of the AlphaFold-derived model (in black cartoon) onto the cryoEM-derived TrwG/VirB8_peri_-TrwE/VirB10_Arches_ structure (in yellow and green cartoon, respectively). RMSD is 0.65 Å. **C.** Top view of the structure of the asymmetric unit of the Arches. Left: the four TrwG/VirB8 periplasmic and connector domains are shown in cartoon representation, coloured in four shades of yellow, while the poly-Ala chain of the two TrwE/VirB10_Arches_ are shown in green with a surface representation. A dashed rectangle indicates the zoomed-in area shown at right. Right: Zoomed-in view of inset shown at left. This view highlights 1-the 2-fold symmetry between the two TrwG/VirB8_peri-_TrwE/VirB10_Arches_ complexes; 2-the involvement of TrwG/VirB8_peri_ α4 and α5 in binding TrwE/VirB10_Arches_. **D.** Details of secondary structures participating in TrwG/VirB8_peri_-TrwE/VirB10_Arches_ interaction. Both proteins are as in panel C, except for the TrwG/VirB8_peri_ region shown in red which points to a region of VirB8_peri_ shown previously to interact with VirB10 (Sharifahmadian *et al*, 2017). Secondary structures in TrwG/VirB8_peri_ are labelled, showing interaction in TrwG/VirB8 is principally along the α4 and α5 helices. Residues boundaries for TrwE/VirB10_Arches_ are labelled. Arrows indicate which T4SS sub-complex to which TrwE/VirB10_Arches_ connect, OMCC at the top, IMC at the bottom.

Because of the two-fold symmetrical arrangement of MolB and MolC, their TrwE/VirB10_Arches_-binding sites locate in opposite faces of the Arches ring, one facing outwards and the other inwards (Figure 4C). Since the inner face of the ring surrounds the VirB2 assembly site on VirB6, this may imply that 6 TrwE/VirB10_Arches_ sequences may contact the pilus in the early phase of pilus biogenesis, potentially providing a sensing or checkpoint mechanism for pilus assembly.

### The inner membrane complex (IMC)

From our previous work (Mace *et al*., 2022), the IMC is a hexamer of an IMC protomer which we defined as including two TrwK/VirB4 subunits (TrwK/VirB4_central_ and TrwK/VirB4_outside_), one TrwM/VirB3 subunit, and 3 TrwG/VirB8_tails_. We now define an “extended IMC protomer” containing not only the same set of proteins but also 3 additional polypeptides: the 4^th^ TrwG/VirB8_tail_ mentioned above, the TM region (α1 and α2) of TrwI/VirB6 (residues 30-82), and a newly discovered density, corresponding to TrwE/VirB10 (residues 21 to 69) (in green in Figure 5A). This density runs along TrwI/VirB6 α1 and α2 through the IM (Figure 5B, inset at right), and then extends in the cytoplasm to make limited contacts with the TrwK/VirB4_central_ subunit of one protomer (TrwK/VirB4_central-A_ in Figure 5B, inset at right), followed (as we progress towards its N-terminus) by more extensive contacts with the TrwK/VirB4_central_, TrwM/VirB3, and finally the TrwK/VirB4_outside_ of the adjacent protomer (labelled TrwK/VirB4_central-B_, TrwM/VirB3_-B_ and TrwK/VirB4_outside-B_ in Figure 5C, inset at right). The part of TrwE/VirB10 interacting with the two IM TMs of VirB6 will be referred to as “TrwE/VirB10_IM_” (residues 43 to 69) while the part N-terminal to it interacting with cytoplasm-facing IMC protomer components as “TrwE/VirB10_Cyto_” (residues 21 to 42).

**Figure 5.**
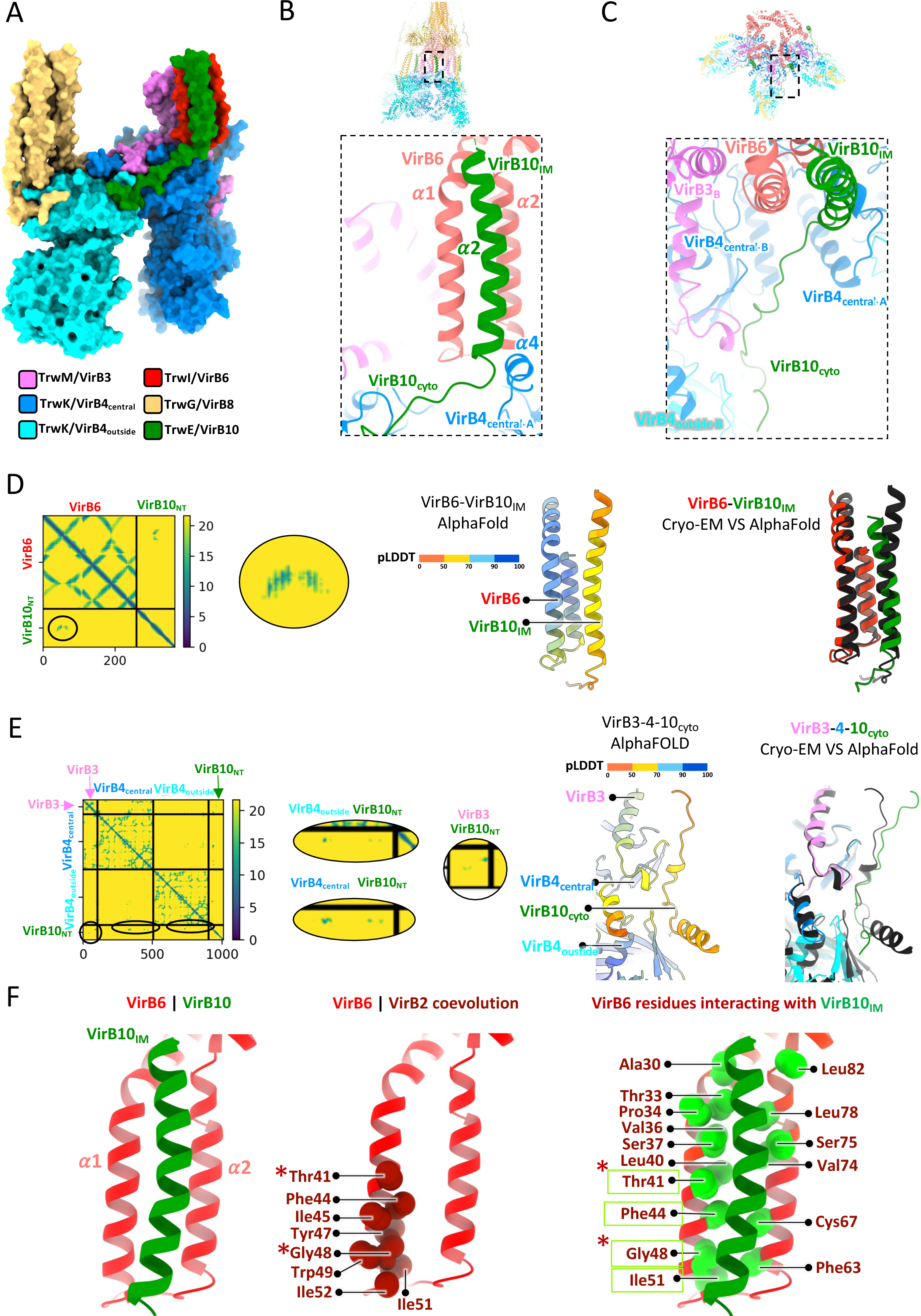
TrwE/VirB10 inner-membrane and cytoplasmic sub-domains. **A.** Structure of the extended IMC protomer. This panel presents a side view of the extended IMC protomer structure in surface representation, color-coded by protein as indicated. The extended IMC protomer includes the formerly-defined IMC protomer (TrwK/VirB4_central_, TrwK/VirB4_outside_, TrwM/VirB3, and four TrwG/VirB8_tails_) to which has been added the TM α1 and α2 of TrwI/VirB6 and TrwE/VirB10_IM_ and TrwE/VirB10_Cyto_. **B.** Location and interaction details of TrwE/VirB10_IM_ with the TM helices (α1 and α2) of TrwI/VirB6. Representation and colour-coding of proteins are as in Figure 1C. Inset locates region of the structure zoomed in at right. **C.** Location and interaction details of TrwE/VirB10_Cyto_ with the TrwK/VirB4_central_-TrwK/VIrB4_outside_-TrwM/VirB3 part of the T4SS complex. Representation and colour-coding of proteins are as in Figure 1C. Inset locates region of the structure zoomed in at right. **D.** Co-evolution of VirB10 with VirB6 and comparison between the cryo-EM and AlphaFold structures of the TrwE/VirB10_IM_-TrwI/VirB6 complex. Left: co-evolution analysis between VirB10_NT_ (see definition of VirB10_NT_ in main text) and VirB6 with co-evolving residue pairs shown as green dots surrounded by a solid-lined oval. A zoom-up of this region is shown at right. Middle: AlphaFold-derived structural model of the TrwE/VirB10_IM_-TrwI/VirB6 complex shown in cartoon representation, colour-coded by model quality (pLDDT). Right: superposition of the AlphaFold-derived (in black cartoon) and cryoEM-derived structure of the TrwE/VirB10_IM_-VirB6_α1α2TM_ complex (in green and red cartoon, respectively). RMSD is 1.067 Å. **E.** Co-evolution of VirB10 with VirB3 and VirB4 and comparison between the cryo-EM and AlphaFold structure of TrwE/VirB10_Cyto_-TrwK/VirB4-TrwM/VirB3 complex. Left: co-evolution analysis between VirB10 N-terminus (VirB10_NT_; see definition of VirB10_NT_ in main text), VirB3 and VirB4, with co-evolving residue pairs involving TrwE/VirB10_NT_ shown here as green dots surrounded by a solid-lined oval. A zoom-up of the various regions of interest is shown at right. Middle: Alphafold-derived structural model of the TrwE/VirB10_Cyto_-TrwK/VirB4-TrwM/VirB3 complex, shown in cartoon representation, colour-coded by model quality (pLDDT). Right: superposition of the AlphaFold-derived (in black cartoon) and cryoEM-derived structures of the TrwE/VirB10_Cyto_-TrwK/VirB4-TrwM/VirB3 complex (in dark and cyan blue cartoon for TrwK/VirB4_central_ and TrwK/VirB4_outside_, respectively, pink cartoon for TrwM/VirB3 and dark green cartoon for TrwE/VirB10_Cyto_). RMSD is 1.85 Å. **F.** TrwI/VirB6 α1 and α2 TM interactions with TrwE/VirB10_IM_. Left: structure of the VirB6 α1 and α2 TM interaction with VirB10_IM_ in cartoon representation coloured in red and green, respectively. Middle: TrwI/VirB6 α1 and α2 in cartoon representation except for their residues known to co-evolve with the VirB2 pilus subunit shown as red spheres. Co-evolved residues are labelled, with a star indicating that the residue was mutated in our previous study and the mutation shown to result in a significant reduction of T4SS conjugation activities. Right: the same view as in left panel, with the cα of TrwI/VirB6 residues interacting with TrwE/VirB10_IM_ shown as green balls and labelled. A green box surrounding a residue name indicates that the VirB6 residue co-evolved in both interactions, with VirB2 and VirB10. Star is as in middle panel.

In these regions of VirB10, the density was of sufficiently high resolution to build side chains. Moreover, AlphaFold predictions of both regions yield models that superimpose well with the model built from the electron density alone (Figure 5, D and E; RMSD of 1.06 and 1.85 Å for TrwE/VirB10_IM_ and TrwE/VirB10_cyto_, respectively). Co-evolution data between VirB10 versus VirB3, VirB4 or VirB6 protein families also confirm the interaction regions (Figure 5, D and E). The first 20 amino acids of TrwE/VirB10 remain invisible, possibly due to disorder or high flexibility.

The interaction between TrwE/VirB10_IM_ with TrwI/VirB6 α1 and α2 are particularly interesting because this binding site for VirB10 on VirB6 entirely overlaps with that of VirB2, the pilus subunit. Indeed, in our earlier work, we identified the site of VirB2-binding on VirB6 using co-evolution analysis and site-directed mutagenesis. We defined this site as “the VirB2 recruitment site” (Mace *et al*., 2022). The co-evolving residues between VirB2 and VirB6 protein families are shown in Figure 5F, 2^d^ panel, while the TrwI/VirB6 residues interacting with TrwE/VirB10_IM_ are shown in Figure 5F, 3^rd^ panel (right-most). As can be seen, the VirB10_IM_-binding site on VirB6 encompasses entirely the VirB2-binding site and therefore VirB10_IM_ in this conformation would prevent VirB2 subunits from being recruited to the VirB6 assembly platform. Therefore, we propose that the regulation of pilus biogenesis is controlled at least in part by the interaction between VirB10_IM_ and the TM region of VirB6. This likely explains our inability to observe a pilus in our biochemical and EM work. The structure of the T4SS we have solved therefore represents an early, inhibited, assembly state preceding pilus biogenesis. We have recently shown that the presence of recipient cell considerably increases pilus biogenesis in the R388 system (AK Vaddakepat, Ho, Waksman, unpublished). We therefore further hypothesize that contacts between donor and recipient cells may be required to release VirB10_IM_ from VirB6, thereby freeing the VirB2 recruitment site and allowing pilus biogenesis to proceed.

From the work presented here emerges a more detailed description of the VirB10 protein. Two sequences have been described earlier: one termed VirB10_O-layer_ (residues 177-395; also known as VirB10_CTD_ (see below)) which together with VirB9_CTD_ and full-length VirB7 forms the O-layer and also forms the T4SS channel though the OM, and another (residue 135-153) referred to as “VirB10_I-layer_“ which forms an α-helix that interacts with VirB9_NTD_ (Chandran *et al*, 2009; Mace *et al*., 2022; Sgro *et al*, 2018). Here, we identify three additional stretches of sequence, VirB10_Arches_ (residues 83-101) that interact with the VirB8_peri_, VirB10_IM_ (residues 43-69) which makes interactions with pilus assembly platform VirB6, and VirB10_cyto_ (residues 21-42) which makes contact with the IMC ATPase complex. Overall, about 72% of the VirB10 sequence is now characterised structurally. A previously-noted sequence (residue 101 to 135) include a proline-rich sequence located between VirB10_Arches_ and VirB10_I-layer_, the function of which remains unclear (Jakubowski *et al*, 2009). The positions, boundary residue numbers, and interaction for all regions are summarized in Figure 6A. One additional fact emerging from this and earlier studies is that VirB10 only contains one folded domain, VirB10_CTD_, functionally overlapping with VirB10_O-layer_ (Figure 6A). The sequence N-terminal to this domain (residue 1-177) has been often referred to as VirB10_NTD_. However, it does not contain a folded domain and therefore it is more accurately named as VirB10 N-terminal sequence or VirB10_NT_ (Figure 6A). These new insights into VirB10 structure and function provides the means to draw a more complete topology diagram for this protein. It is shown in Figure 6B, left panel. It provides an update on the naming of the various secondary structures that VirB10 sequences adopt from the N-terminus in the cytoplasm to the C-terminus near the OM.

**Figure 6.**
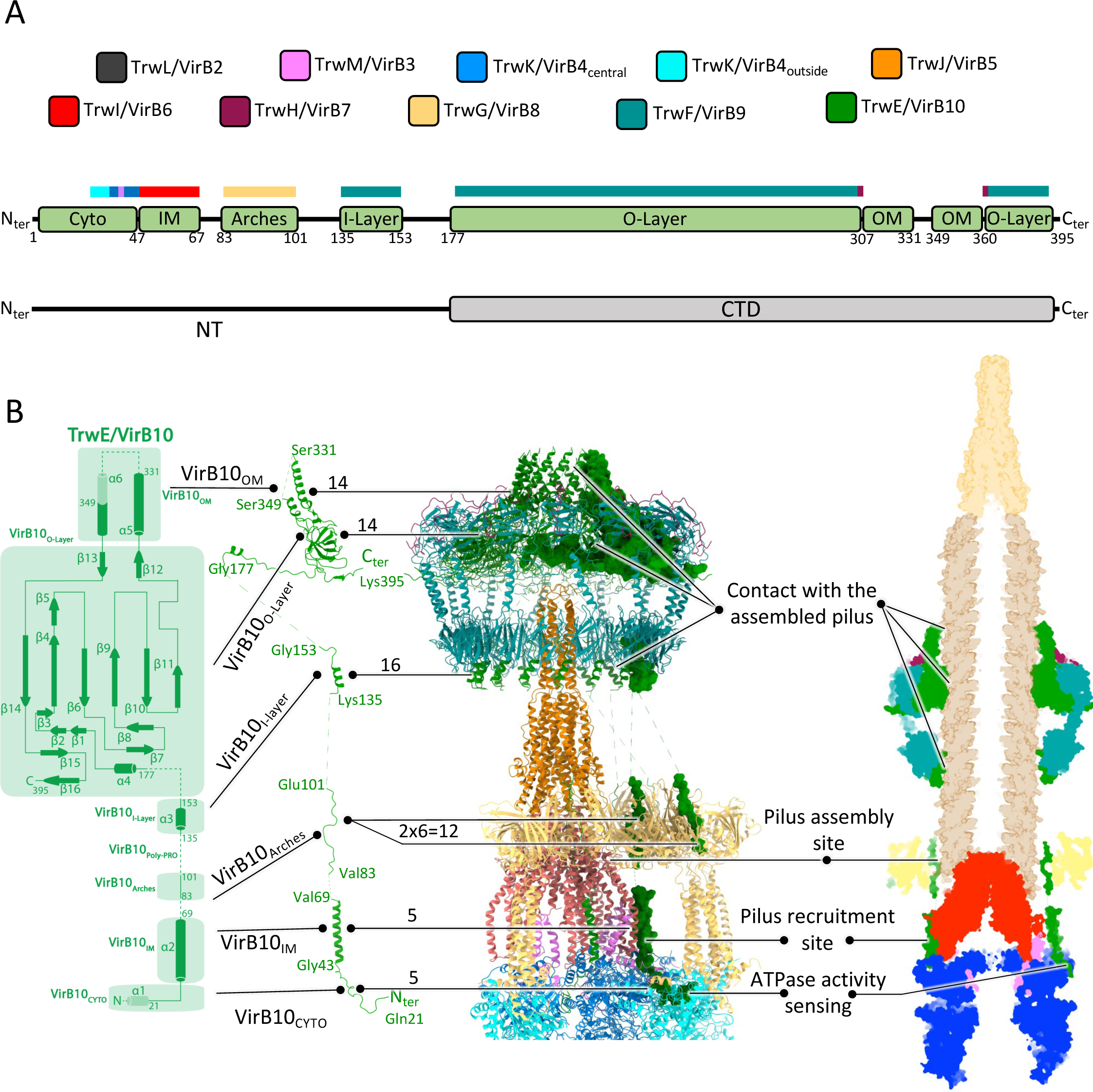
VirB10: a central protein of the T4SS. **A.** Structural organisation and interactions of TrwE/VirB10. Top panel, colour-code for Trw/VirB proteins used in this figure. Middle panel: functional organisation of TrwE/VirB10. Green boxes indicate location and boundary residues of the various functional regions of TrwE/VirB10. In the main text, these functional regions are labelled TrwE/VirB10_XXX_ (for example TrwE/VirB10_Arches_ or TrwE/VirB10_IM_) where the XXX subscript reflects the function and location of that region. The Trw/VirB proteins with which region TrwE/VirB10_XXX_ interact are indicated on the coloured boxes just above (colour is by proteins as in top panel). Bottom panel: folded domain structure of TrwE/VirB10. While two domains were previously described, only one is now known to adopt a defined fold, TrwE/VirB10_CTD_, which is part of the O-layer et forms the OM channel. The previously named NTD is a linear peptide best described as an N-terminal region or TrwE/VirB10_NT_. **B.** TrwE/VirB10 full-length structure details. Left: a topology diagram of TrwE/VirB10, with annotated domains and secondary structures. Middle left: The full-length VirB10 structure is presented in cartoon representation. Middle right: The T4SS structure is shown in cartoon representation, except for TrwE/VirB10, which is displayed in surface representation. The number of VirB10 copies known to make interactions with other VirB proteins or itself in the various T4SS regions and sub-complexes is indicated. Right: A cut-out side view in surface representation of the T4SS-pilus model, with the pilus and its TrwJ/VirB5 tips. The VirB10’s potential functions by region/domain during pilus biogenesis are indicated. See main text for details.

Zooming out to obtain an overall view of the VirB10 protein within the T4SS structure as shown in Figure 6B, right panels, it becomes clear that VirB10 can potentially make contact with the pilus all along the pilus length and with its assembly site, suggesting it may play a major role in regulating and/or facilitating pilus biogenesis at many different stages. Not only the two channel-forming α-helices (termed VirB10_OM_ in Figures 6) would make contact with the pilus as it is threaded through the OM channel, but also a residue of the periplasmic part of VirB10_O-layer_ has been shown to be potentially involved in the gating mechanism of that channel (Banta *et al*, 2011; Chandran *et al*., 2009). Further downstream, VirB10_I-layer_ could potentially contact the pilus. One of the two VirB10_Arches_ that we have observed runs very close to the VirB2 assembly site on VirB6 and we now know that VirB10_IM_ in the IM obstructs the VirB2 recruitment site on VirB6, providing two other means by which VirB10 may be regulating pilus subunit recruitment and assembly. Finally, we observe a sequence of VirB10 (VirB10_Cyto_) that makes multiple contact with the TrwK/VirB4 ATPase, thereby potentially providing another point of pilus biogenesis regulation.

An intriguing feature of the T4SS structure solved previously was the paucity of interactions between the various sub-complexes (OMCC, Arches, Stalk and IMC). However, from the work presented here (Figure 6B), VirB10 emerges as a crucial and unique element that “glues” these distinct subcomplexes together.

Another remarkable feature of VirB10 is that the T4SS include 16 copies of them, only 14 are used in the O-layer, potentially only 12 of them in the Arches, with only 5 of those involved in VirB6 binding and 6 involved in VirB4 ATPase contacts (Figure 6, middle panel). Understanding such a striking mismatch symmetry and why so many VirB10 are needed when so few may be used is one of the most puzzling features of the T4SS, a puzzle that further structural work will undoubtedly solve.

## CONCLUSION

The study presented here provides additional insights that shed new lights onto several aspects of T4SS structure and function. We were able to provide further details on the structural dynamics of the OMCC and the various conformations it can adopt. We characterised a hydrophobic tip at the N-terminus of VirB5, and a 4^th^ VirB8 subunits adding to the Arches and the IMC. However, the functional significance of these new structural features remains unclear. More functionally significant perhaps has been the characterisation of a VirB10 region that has the potential to interfere with an essential T4SS function which is involved in pilus biogenesis. Indeed, VirB10 proteins define a rather rare class of membrane proteins that spans both the IM and OM of Gram-negative bacteria. Using two-hybrid or phage-display methods as well as biochemical screening of peptide libraries, VirB10 has been shown to interact with most T4SS components (Mary *et al*, 2018; Sharifahmadian *et al*., 2017; Terradot *et al*, 2004). One well-documented interaction of VirB10 is with the VirD4 ATPase, the so-called coupling protein because it couples substrate recruitment to substrate transfer (Llosa *et al*, 2003). However, the molecular basis for any of these roles has been unclear. Here, in addition to further details concerning the structure of the entire T4SS, we reveal a new role for VirB10 in pilus biogenesis and provide the molecular basis for this role. Given the considerable importance that the pilus plays in conjugation, unravelling novel regulatory mechanisms for its biogenesis should provide new approaches targeting their inhibition, potentially leading to the design of new tools to stop the spread of antibiotic resistance genes.

## MATERIALS and METHODS

### Bacterial strains, constructs, expression and purification of T4SS

R388 T4SS complexes were expressed and purified from membranes as described previously (Mace *et al*., 2022).

### Cryo-EM grid preparation and data acquisition

Grids preparation and data acquisition were as described previously (Mace *et al*., 2022).

### EM processing

MOTIONCOR2 (Zheng *et al*, 2017) was employed for motion correction and dose weighting, followed by CTF estimation using CTFFIND v4.1 (Rohou & Grigorieff, 2015). EM processing workflows for the various parts of the T4SS are shown in Figures EV1 and EV2.

### Image processing of the T4SS OMCC

#### Pre-processing

Reprojections of a low pass filtered (20 Å) map generated using PDB 3JQO (Chandran *et al*., 2009) were used to pick particles centred on the OMCC with GAUTOMATCH v0.56 (Zhang, 2017). Following multiple rounds of 2D classification, 1,742,107 particles were selected. After 3D heterogeneous classification without symmetry applied, the two best classes were selected, resulting in a selection of 1,360,271 particles that will be used in all subsequent processing. A refined map at 3.12 Å resolution without symmetry applied was then obtained, that served as reference map in all subsequent processing.

#### O-layer and I-layer high resolution

Reference map and particles were imported into RELION 3.1 (Scheres, 2012) for 3D classification without alignment. The best class was used in two rounds of local refinements using CRYOSPARC (Punjani *et al*, 2017), one focused on the O-layer with C14 symmetry applied, and the other comprising the I-Layer with C16 symmetry applied. The resulting electron density maps had an average resolution of 2.46 Å for the O-Layer (map termed “O-layer C14 at 2.5 Å” in Table EV1) and 2.69 Å for the I-Layer (map termed “I-layer C16 at 2.7 Å” in Table EV1) as estimated using the gold standard Fourier Shell Correlation (FSC) with a 0.143 threshold.

#### OMCC heterogenicity analysis

Reference map and particles were used for heterogeneous refinement without symmetry applied into five classes. After visual inspection in CHIMERA v1.4 (Pettersen *et al*, 2004), particles from classes showing the same I-layer conformation were combined for a final homogeneous refinement using CRYOSPARC, resulting in three maps: Conformation-A, -B and -C at resolutions of 3,18 Å, 2.93 Å and 3.05 Å, respectively (Table EV1.

### Image processing of the T4SS Stalk-Arches-IMC

#### Pre-processing

Reprojections of the negative-strain EM map of the IMC, Stalk and Arches (EMDB 3585 (Redzej *et al*, 2017)) were used to pick particles centred on the IMC, Stalk and Arches using GAUTOMATCH. Following multiple rounds of 2D classification using CRYOSPARC, 1,0482,424 particles were selected. After 3D heterogeneous classification without symmetry applied using CRYOSPARC, the two best classes were selected, resulting in a selection of 600,997 particles and a refinement map at 7.35 Å resolution without symmetry being applied. Both particles and map were used as reference for subsequent analysis.

#### Stalk analysis

From the reference map and particles, two rounds of local 3D classification (using the Stalk mask described previously (Mace *et al*., 2022)) were performed using RELION. The final best class, composed of 104,720 particles was selected for a round of Non-Uniform refinement (NU-refinement) with C5 symmetry applied using CRYOSPARC, yielding to a 2.97 Å resolution map (termed “Stalk C5 at 3.0 Å” in Table EV1).

#### Arches analysis

From the reference map and particles, a round of local 3D classification was performed using RELION and a previously described mask (Mace *et al*., 2022). The best class, composed of 115,034 particles, was selected to perform a round of NU-refinement without symmetry applied using CRYOSPARC, yielding a 6.22 Å resolution map (termed “Arches at 6.2 Å” in Table EV1).

#### Extended IMC protomer analysis

From the reference map and particles, a round of local 3D classification was performed using RELION and a mask similar to that described in our previous study for the IMC protomer (defined at the time to only contains two TrwK/VirB4, one TrwM/VirB3, and 4 TrwG/VirB8_tails_) but, this time, extended to encompass the TrwE/VirB10 IM and Cyto regions, the TrwI/VirB6 TM region (α1 and α2) and the non-interpretable density shown in Figure 1, box 7 (Figure EV2). The best class, composed of 234,578 particles was selected to perform NU-refinement without symmetry, applied using CRYOSPARC, yielding to a 3.83 Å resolution map (termed “extended IMC protomer at 3.8 Å” in Tables 1 and S1).

#### Stalk-Arches-IMC analysis

From the reference map and particles, two local 3D classifications were performed using RELION and a mask surrounding the area of interest. The final best class, composed of 65,178 particles, was selected to perform NU-refinement without symmetry applied using RELION, yielding a 4.33 Å resolution map (map termed “Stalk-Arches-IMC at 4.3 Å”).

All maps were subjected to sharpening using DEEPEMHANCER (Sanchez-Garcia *et al*, 2021) in “tightTarget” mode, and local resolution estimated using CRYOSPARC. Detailed map statistics are provided in Table EV1.

### Structures model building

#### For the OMCC

We employed the previously-determined O-layer (PDB-7O3J (Mace *et al*., 2022)) and I-layer (PDB-7O3T (Mace *et al*., 2022)) R388 structures as starting models. These models were manually fitted into the new O-layer and I-layer highest resolution maps (those where either C14 or C18 symmetry was applied, respectively (see above)) using COOT v0.9.3 (Emsley & Cowtan, 2004) and refined using PHENIX v1.18.2 (Adams *et al*, 2010) with secondary structural elements and Ramachandran restraints applied. To generate the OMCC conformations A, B, and C models, the O- and I-layer structures were docked into the O- and I-layer densities of the corresponding maps and the linker helix between TrwF/VirB9 CTD and NTD was built and fitted manually (using COOT) into density for all TrwF/VirB9 subunits where the density if visible (14 subunits out of 16). The resulting models were refined using PHENIX, ultimately yielding a final model for Conformation-A, -B and -C OMCC structures (Table EV1). *<u>For the Stalk</u>*. We used the previously solved Stalk structure (PDB-7O3V (Mace *et al*., 2022) as a starting model to build a more complete model into the higher resolution map that we describe here (“Stalk C5 at 3.0 Å” map in Table EV1) using COOT. This structure incorporates new elements, such as the TrwJ/VirB5 tip and TrwI/VirB6 transmembrane domains. This model was then refined using PHENIX to generate the final Stalk structure.

#### For the Arches

We started with the previously solved Arches structure (PDB-7OIU (Mace *et al*., 2022)). To this structure, using the “Arches at 6.2 Å” map, we added in the asymmetric unit of the Arches a 4^th^ TrwG/VirB8_peri_ domain and the TrwG/VirB8 connector domain derived from AlphaFold modelling. All the sidechains were removed from this model, and further improvements were manually achieved using COOT. PHENIX was used for final refinement (Table EV1).

#### For the extended IMC protomer

We began with the previously solved IMC protomer structure (PDB-7Q1V (Mace *et al*., 2022)). To this structure and using the “extended IMC protomer at 3.8 Å” map, we built a 4^th^ TrwG/VirB8_tail_, the TrwI/VirB6_TM_ helices (α1-α2), and the VirB10_IM_ and VirB10_cyto_ regions. This initial model underwent manual improvements using COOT and final refinement using PHENIX, leading to the extended IMC protomer structure containing not only the two TrwK/VirB4 subunits, the TrwM/VirB3 subunit and the 4 TrwG/VirB8_tails_, but also the TrwI/VirB6 TM region as well as the TrwE/VirB10 IM and cyto regions (Table EV1).

#### For the Stalk-Arches-IMC

The structural model was constructed using the Stalk-Arches-IMC at 4.3 Å map and the high-resolution solved structures of the Stalk, Arches asymmetric unit and extended IMC protomer. This map was important to characterise contacts of TrwE/VirB10_CYTO_ that are shared with two adjacent TrwK/VirB4 subunits as shown in Figure 5C, right panel. In this composite model, regions with poor Cα backbone density were removed as well as all side chains. COOT was employed for manual improvement, and PHENIX was used for final refinement to generate the Stalk-Arches-IMC structure (Table EV1).

Quality assessments of all structures were performed using MolProbity v4.5.1 (Davis *et al*, 2007). Data and model statistics are provided in Table S1B.

EM maps and atomic models were deposited to the EMDB and PDB data bases. Accession codes can be found in SI Table 1 of the manuscript.

### Structure validation using co-evolution analysis and AlphaFold structural prediction

To validate the new structural features and interactions presented in this study, we utilised AlphaFold (Jumper *et al*., 2021) through the ColabFold advanced notebook (Mirdita *et al*, 2022). This approach allowed us to generate models and co-evolution diagrams as structural validation tools of our EM-derived models.

### Interaction analysis and representation of the T4SS structure

Interaction analysis was carried out using the PISA server (Krissinel & Henrick, 2007), and structure figures were generated using ChimeraX v1.6 (Pettersen *et al*, 2021).

## ACKNOWLEDGEMENTS

We would like to thank Dr. David Houldershaw for IT support and Natasha Lukoyanova for preparation of EM grids and collection of data. This work was supported by Wellcome grants 098302 and 217089 to GW. Cryo-EM data for this investigation were collected at the ISMB EM facility at Birkbeck College, University of London with financial support from Wellcome Trust (202679/Z/16/Z and 206166/Z/17/Z). KM purified the T4SS complex. EM processing and AlphaFOLD analysis work was carried out by KM and KM built and refined the structure. KM and GW wrote the manuscript. GW supervised the work.

## REFERENCES

Adams PD, Afonine PV, Bunkoczi G, Chen VB, Davis IW, Echols N, Headd JJ, Hung LW, Kapral GJ, Grosse-Kunstleve RW et al (2010) PHENIX: a comprehensive Python-based system for macromolecular structure solution. Acta Crystallogr D Biol Crystallogr 66: 213–221

Aly KA, Baron C (2007) The VirB5 protein localizes to the T-pilus tips in *Agrobacterium tumefaciens*. Microbiology 153: 3766–3775

Amin H, Ilangovan A, Costa TRD (2021) Architecture of the outer-membrane core complex from a conjugative type IV secretion system. Nat Commun 12: 6834

Banta LM, Kerr JE, Cascales E, Giuliano ME, Bailey ME, McKay C, Chandran V, Waksman G, Christie PJ (2011) An *Agrobacterium* VirB10 mutation conferring a type IV secretion system gating defect. J Bacteriol 193: 2566–2574

Barlow M (2009) What antimicrobial resistance has taught us about horizontal gene transfer. Methods Mol Biol 532: 397–411

Chandran Darbari V, Waksman G (2015) Structural Biology of Bacterial Type IV Secretion Systems. Annu Rev Biochem 84: 603–629

Chandran V, Fronzes R, Duquerroy S, Cronin N, Navaza J, Waksman G (2009) Structure of the outer membrane complex of a type IV secretion system. Nature 462: 1011–1015

Costa TRD, Harb L, Khara P, Zeng L, Hu B, Christie PJ (2021) Type IV secretion systems: Advances in structure, function, and activation. Mol Microbiol. 115: 436–452

Costa TRD, Ilangovan A, Ukleja M, Redzej A, Santini JM, Smith TK, Egelman EH, Waksman G (2016) Structure of the Bacterial Sex F Pilus Reveals an Assembly of a Stoichiometric Protein-Phospholipid Complex. Cell 166: 1436–1444 e1410

Costa TRD, Patkowski JB, Mace K, Christie PJ, Waksman G (2023) Structural and functional diversity of type IV secretion systems. Nat Rev Microbiol. doi: 10.1038/s41579-023-00974-3.

Davis IW, Leaver-Fay A, Chen VB, Block JN, Kapral GJ, Wang X, Murray LW, Arendall WB, 3rd, Snoeyink J, Richardson JS et al (2007) MolProbity: all-atom contacts and structure validation for proteins and nucleic acids. Nucleic Acids Res 35: W375–383

Durie CL, Sheedlo MJ, Chung JM, Byrne BG, Su M, Knight T, Swanson M, Lacy DB, Ohi MD (2020) Structural analysis of the *Legionella pneumophila* Dot/Icm type IV secretion system core complex. Elife 9:e59530

Emsley P, Cowtan K (2004) Coot: model-building tools for molecular graphics. Acta Crystallogr D Biol Crystallogr 60: 2126–2132

Jakubowski SJ, Kerr JE, Garza I, Krishnamoorthy V, Bayliss R, Waksman G, Christie PJ (2009) *Agrobacterium* VirB10 domain requirements for type IV secretion and T pilus biogenesis. Mol Microbiol 71: 779–794

Jumper J, Evans R, Pritzel A, Green T, Figurnov M, Ronneberger O, Tunyasuvunakool K, Bates R, Zidek A, Potapenko A et al (2021) Highly accurate protein structure prediction with AlphaFold. Nature 596: 583–589

Krissinel E, Henrick K (2007) Inference of macromolecular assemblies from crystalline state. J Mol Biol 372: 774–797

Llosa M, Zunzunegui S, de la Cruz F (2003) Conjugative coupling proteins interact with cognate and heterologous VirB10-like proteins while exhibiting specificity for cognate relaxosomes. Proc Natl Acad Sci U S A 100: 10465–10470

Mace K, Vadakkepat AK, Redzej A, Lukoyanova N, Oomen C, Braun N, Ukleja M, Lu F, Costa TRD, Orlova EV et al (2022) Cryo-EM structure of a type IV secretion system. Nature 607: 191–196

Mary C, Fouillen A, Bessette B, Nanci A, Baron C (2018) Interaction via the N terminus of the type IV secretion system (T4SS) protein VirB6 with VirB10 is required for VirB2 and VirB5 incorporation into T-pili and for T4SS function. J Biol Chem 293: 13415–13426

Mirdita M, Schutze K, Moriwaki Y, Heo L, Ovchinnikov S, Steinegger M (2022) ColabFold: making protein folding accessible to all. Nat Methods 19: 679–682

Pettersen EF, Goddard TD, Huang CC, Couch GS, Greenblatt DM, Meng EC, Ferrin TE (2004) UCSF Chimera--a visualization system for exploratory research and analysis. J Comput Chem 25: 1605–1612

Pettersen EF, Goddard TD, Huang CC, Meng EC, Couch GS, Croll TI, Morris JH, Ferrin TE (2021) UCSF ChimeraX: Structure visualization for researchers, educators, and developers. Protein Sci 30: 70–82

Punjani A, Rubinstein JL, Fleet DJ, Brubaker MA (2017) cryoSPARC: algorithms for rapid unsupervised cryo-EM structure determination. Nat Methods 14: 290–296

Redzej A, Ukleja M, Connery S, Trokter M, Felisberto-Rodrigues C, Cryar A, Thalassinos K, Hayward RD, Orlova EV, Waksman G (2017) Structure of a VirD4 coupling protein bound to a VirB type IV secretion machinery. EMBO J 36: 3080–3095

Rohou A, Grigorieff N (2015) CTFFIND4: Fast and accurate defocus estimation from electron micrographs. J Struct Biol 192: 216–221

Sanchez-Garcia R, Gomez-Blanco J, Cuervo A, Carazo JM, Sorzano COS, Vargas J (2021) DeepEMhancer: a deep learning solution for cryo-EM volume post-processing. Commun Biol 4: 874

Scheres SH (2012) RELION: implementation of a Bayesian approach to cryo-EM structure determination. J Struct Biol 180: 519–530

Sgro GS, Costa TR, Cenens W, Souza DP, Cassagno A, Oliveira LC, Salinas RK, Portugal RV, Farah CS, Waksman G (2018) Cryo-EM structure of the core complex of a bacterial killing type IV secretion system. Nature Microbiology. 3: 1429–1440

Sharifahmadian M, Nlend IU, Lecoq L, Omichinski JG, Baron C (2017) The type IV secretion system core component VirB8 interacts via the beta1-strand with VirB10. FEBS Lett 591: 2491–2500

Sheedlo MJ, Chung JM, Sawhney N, Durie CL, Cover TL, Ohi MD, Lacy DB (2020) Cryo-EM reveals species-specific components within the Helicobacter pylori Cag type IV secretion system core complex. Elife 9:e59495.

Terradot L, Bayliss R, Oomen C, Leonard GA, Baron C, Waksman G (2005) Structures of two core subunits of the bacterial type IV secretion system, VirB8 from *Brucella suis* and ComB10 from Helicobacter pylori. Proc Natl Acad Sci U S A 102: 4596–4601

Terradot L, Durnell N, Li M, Li M, Ory J, Labigne A, Legrain P, Colland F, Waksman G (2004) Biochemical characterization of protein complexes from the *Helicobacter pylori* protein interaction map: strategies for complex formation and evidence for novel interactions within type IV secretion systems. Mol Cell Proteomics 3: 809–819

Virolle C, Goldlust K, Djermoun S, Bigot S, Lesterlin C (2020) Plasmid Transfer by Conjugation in Gram-Negative Bacteria: From the Cellular to the Community Level. Genes (Basel) 11:1239

Waksman G (2019) From conjugation to T4S systems in Gram-negative bacteria: a mechanistic biology perspective. EMBO Rep 20:e47012

Wu X, Zhao Y, Sun L, Jiang M, Wang Q, Wang Q, Yang W, Wu Y (2019) Crystal structure of CagV, the *Helicobacter pylori* homologue of the T4SS protein VirB8. FEBS J 286: 4294–4309

Zhang K (2017) Fully automatic acccurate, convenient and extremely fast particle picking for EM. https://www.mrc-lmbcamacuk/kzhang/Gautomatch/

Zheng SQ, Palovcak E, Armache JP, Verba KA, Cheng Y, Agard DA (2017) MotionCor2: anisotropic correction of beam-induced motion for improved cryo-electron microscopy. Nat Methods 14: 331–332

